# Host traits drive viral life histories across phytoplankton viruses

**DOI:** 10.1101/124743

**Authors:** Kyle F. Edwards, Grieg F. Steward

## Abstract

Viruses are integral to ecological and evolutionary processes, but we have a poor understanding of what drives variation in key traits across diverse viruses. For lytic viruses, burst size, latent period, and genome size are primary characteristics controlling host-virus dynamics. Burst size and latent period are analogous to organismal traits of fecundity and generation time, and genome size affects the size of the virion as well as viral control of host metabolism. Here we synthesize data on these traits for 75 strains of phytoplankton viruses, which play an important role in global biogeochemistry. We find that primary traits of the host (genome size, growth rate) are major ecological drivers, explaining 40-50% of variation in burst size and latent period. We analyze an eco-evolutionary model to explore mechanisms underlying these patterns. We find that burst size may be set by the host genomic resources available for viral construction, while latent period evolves to permit this maximal burst size, modulated by host metabolic rate. These results suggest that general mechanisms may underlie the evolution of diverse viruses, which will facilitate our understanding of viral community processes, ecosystem impacts, and coevolutionary dynamics.

## Introduction

Viruses are integral to the ecology and evolution of all cellular life (Villarreal and Witzany 2010, Koonin and Dolja 2013). By selectively lysing or altering the physiology of the cells they infect, and mediating horizontal gene transfer among genetically distinct lineages, they can generate and maintain host diversity and influence biogeochemical cycles (Suttle 2007, Breitbart 2012). Viruses are highly host-specific and exhibit great taxonomic and genetic diversity, with many forms still being discovered and characterized (Lang et al. 2009, Rosario et al. 2012, Fischer 2016). Although viruses were discovered over a century ago, and some model systems have been studied in great detail, we have a poor understanding of what drives variation in key traits across diverse viruses. Research on functional trait diversity has been fruitful in linking physiology, community structure, and ecosystem dynamics, particularly in terrestrial plants (Westoby and Wright 2006) and phytoplankton (Litchman and Klausmeier 2008). If broad patterns and mechanisms underlying how viruses function can be discerned, this will promote a general framework for predicting viral ecology and the effects of viral interactions on host populations, communities, and ecosystems (Gudelj et al. 2010). In this study we examine viruses that infect phytoplankton, a group of viruses that have a direct influence one of the most important biogeochemical transformations on the planet. Phytoplankton account for nearly half of global primary production and participate in multiple elemental cycles (Falkowski et al. 2004). Viral effects on mortality, element cycling, and community structure of phytoplankton potentially have global consequences (Brussaard 2004, Weitz et al. 2015).

We will focus on several traits of lytic viruses, which always produce virions and lyse the host cell, provided the cell has sufficient resources and integrity to support the infection. Viruses replicating via the lytic cycle can have particularly dramatic effects on host populations, because of their rapid rate of reproduction and corresponding destruction of the host population. Among the key traits that define the function of lytic viruses are burst size (new virions produced per infected host), latent period (time elapsed between infection and lysis), and viral genome size. Burst size and latent period are analogous to the organismal life history traits of fecundity and generation time, and are key parameters for host-virus population dynamics. The burst size and latent period of a virus can vary with environmental conditions or host genotype (Wilson et al. 1996, Maat and Brussaard 2016), but these parameters also vary greatly among different viral strains measured under similar conditions (typically, resource-replete exponential growth of the host). For simplicity we will refer to differences across isolates as variation in ‘viral traits’, although such differences are likely driven by both viral genotype and host genotype. Viral genome size is a trait that will influence the ability of the virus to control host metabolism, while also determining the metabolic cost of synthesizing new virions (Bragg and Chisholm 2008, Thomas et al. 2011), as well as the size of the virion (Steward et al. 2013), which will affect diffusivity and other processes (Murray and Jackson 1992). Here we analyze variation in burst size, latent period, and genome size across viruses, because these traits have been quantified for numerous isolates that infect phytoplankton and other unicellular algae.

Guided by previous work, we can make predictions about drivers of variation in these traits: conditions that may select for particular trait values, and constraints that may cause the traits to covary. Viruses use their host’s molecular machinery to reproduce, and therefore host structure and physiology are primary selective forces for viral trait evolution. Nucleotides for viral genome synthesis come from host nucleotide pools and degradation of the host genome, with an uncertain and variable contribution of *de novo* nucleotide synthesis during infection (Van Etten et al. 1984, Wikner et al. 1993, Brown et al. 2007, Thompson et al. 2011). Host genome size may thus influence the rate of viral production or burst size, and a prior synthesis of 15 host-virus pairs found that viral nucleotide production correlates with host genome size (Brown et al. 2006). The growth rate of the host will likely correlate with the concentration of ribosomes and enzymes required to synthesize host proteins, RNA, and DNA, which are also used to construct new virus particles (You et al. 2002, Daines et al. 2014). Therefore, host growth rate may also affect the rate of viral production.

Within the context set by host conditions, viral traits should evolve to maximize fitness. Burst size and latent period are intrinsically related, because lysing the host sooner will reduce burst size, all else equal. Theory and experiments with *E. coli* show that the optimal latent period (and burst size) depends on host density and quality, such that a shorter latent period is selected for when hosts are more dense or of higher quality (Wang et al. 1996, Abedon et al. 2003). However, these models and experiments assume that host density is relatively constant, rather than being regulated by the virus, and so it is less clear how latent period will evolve when host and virus populations are dynamic. Furthermore, the benefit of a longer latent period may be diminished if viral reproduction exhausts nucleotide resources within the host. Models that include bacteriophage-host dynamics and latent period evolution have been developed recently (Bonachela and Levin 2014). By parameterizing an eco-evolutionary model for phytoplankton-virus interactions, and manipulating host traits relevant to our hypotheses, we can test whether empirical patterns are consistent with theoretical expectations for particular mechanisms.

In sum, we can predict that burst size and latent period will tend to be correlated across viral strains, but that both of these traits may be modulated by the genome size of the host (larger hosts may permit a larger optimal burst size), the growth rate of the host (faster growth may decrease optimal latent period), and the genome size of the virus (larger viruses require more resources per virion, which may reduce optimal burst size). To test these predictions we compile published data on burst size, latent period, and genome size for lytic viruses isolated from phytoplankton/microalgae. We explore hypothesized relationships among virus traits, and between virus and host traits. We also analyze a model of viral trait evolution to explore potential mechanisms for the empirical patterns of trait variation.

## Methods

### Virus trait compilation

The literature was searched for studies that measured burst size and latent period of viruses isolated on phytoplankton or other microalgal hosts (Appendix Table S1). We only included experiments where the host was grown under nutrient-replete conditions, so that we could quantify functional variation across viruses cultured under similar conditions, as opposed to plastic responses of individual strains. We recorded the name of the virus strain, virus genome type (dsDNA/dsRNA/ssDNA/ssRNA), virus source location, host species, host taxon (chlorophyte/cryptophyte/cyanobacterium/diatom/dinoflagellate/haptophytepelagophyte/raphidophyte), and environment (marine/freshwater). We also recorded whether burst size was estimated by counting infectious units or free virions. Virus capsid size (diameter) and genome size estimates were taken from the same study, or other studies on the same isolate. Virus genome size correlates strongly with capsid diameter, and thus we only use genome size to represent virus size in all analyses (Fig. S1). For some analyses we quantify total viral nucleotide output at lysis (burst size * virus genome size); for these calculations we divided the genome size of the single-stranded viruses by a factor of 2. Host genome size and cell volume estimates were taken from the literature, and if an estimate for the host species was not available, an estimate from a congener was used if available. It is noteworthy that 10 of the 13 single-stranded viruses have been isolated from *Chaetoceros* species, and, in the absence of published information on genome sizes for most of the hosts, we assigned all of the host species the same genome size (measured for *C. muelleri*). For the double-stranded viruses, genome size estimates were available for nearly all host species. When possible, host exponential growth rate was estimated using DataThief (Tummers 2006) to extract growth curves measured on uninfected hosts, or hosts growing prior to infection. Temperature and irradiance under which the hosts were cultured during one-step growth experiments were also recorded.

### Statistical methods

Relationships between viral traits, or viral traits and host traits, were analyzed using mixed models (R package lme4; Bates et al. 2015). We used random effects to appropriately account for non-independence in the data resulting from multiple viruses infecting similar hosts (host genus), multiple viruses measured in the same study (study ID), host taxonomy (diatom/cyanobacteria/haptophyte/etc.), or similarities among virus type (dsDNA/dsRNA/ssDNA/ssRNA). These random effects were included in all models, and different fixed effects and response variables were used to test different relationships (e.g., host genome size as a predictor of burst size). Significance of fixed effects was tested using approximate F-tests (R package lmerTest; Kuznetsova et al. 2016), and variation explained by fixed effects was quantified as marginal R^2^GLMM (R package MuMIn; Bartón 2016). Preliminary results showed that patterns did not vary between marine and freshwater strains, or between estimates of burst size using infectious units vs. direct counts; therefore these factors were excluded from the analyses for simplicity.

### Model of evolution of latent period / burst size

As shown below, there is evidence that viral traits are affected by the genome size and growth rate of the host. To ask how viral strategies are expected to evolve across phytoplankton hosts that vary in growth rate and genome size, we analyzed a model of viral trait evolution. The model is adapted from previous work on bacteriophage (Levin et al. 1977, Bonachela and Levin 2014), and the contribution of our analysis is to test the effect of parameters representing host growth and genome size. The model can be written as follows:

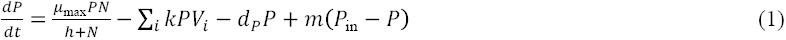

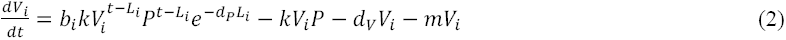

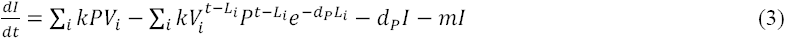

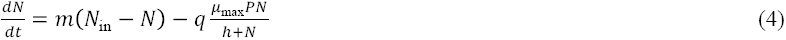

The model explicitly represents latent period as a delay between infection and production of new virions (Bonachela and Levin 2014). In eqn. (1), susceptible phytoplankton (*P*) grow, limited by nutrient (*N*), with maximum growth rate μ_max_ and half-saturation constant *h*. Susceptible hosts are lost due to infection from each viral strain *i* (*V*_i_), with adsorption rate *k*. There is loss due to other mortality *d*_p_, which is meant to primarily represent grazing. There is also a slow rate of mixing with adjacent waters (*m*), which causes susceptible hosts from elsewhere (*P*_in_) to enter the system. In eqn. (2), viral strain *i* has burst size *b*_i_, produced from hosts that were infected at time (*t*-*L*i), where *L*_i_ is the latent period. The term *e^−d_p_L_i_^* represents the fraction of infected hosts that have not died by the end of the latent period. Free virions are lost as a result of adsorption to new hosts, decay at rate *d*_V_, or mixing at rate *m*. In eqn. (3), infected hosts are created through adsorption, are lost to viral lysis, other sources of mortality, or mixing. In eqn. (4), nutrients from elsewhere (*N*_in_) enter the system by mixing, and are taken up during phytoplankton growth, with constant cellular quota *q*. This model does not include recycling of phytoplankton nutrients from lysis or other mortality for computational simplicity; we have checked to ensure this does not affect the trait evolution results.

To explore the influence of host characteristics, viral burst size and latent period are related to each other and to host genome size and growth rate:

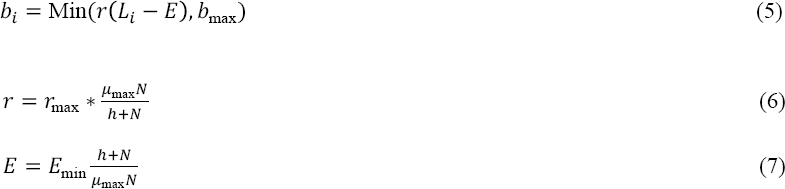

Where *r* is the linear rise rate for viral particle production and *E* is the eclipse period. We assume *r* is proportional to host growth rate (with maximum *r*_max_^*^*μ_max_*), and *E* is inversely proportional to host growth rate (with minimum *E_min_/*μ*_max_*). The functional form of eqns. (5-7) are based on the infection cycle of phage T7 infecting *E. coli* (You et al. 2002). To represent limitation of particle production by host genomic resources, burst size is also given an upper limit *b*_max_, which is meant to represent the ratio of host genome size: virus genome size. The magnitude of this upper limit is varied to test the effect of host genomic resources on viral life history strategy. The magnitude of *μ_max_* is varied to test the effect of host growth rate.

To simulate evolution under this model, 100 virus genotypes were initialized at equal density, with latent periods ranging from 3.5 to 96 hrs, and corresponding burst sizes calculated from the above equations. All virus genotypes compete for a single host, and the model was run until one virus genotype competitively excluded all others. The delay differential equations were solved with LSODA in the R package deSolve (Soetaert et al. 2010). External inputs to the model are constant (it is a nutrient-limited chemostat / mixed layer model), but the host and viruses oscillate substantially in abundance (Fig. S2). The amplitude of the oscillations is somewhat reduced by the diffusion of susceptible hosts into the system, which aids in computational tractability. Parameter values were chosen to represent a typical phytoplankton-virus system (Table 1).

## Results

### Compilation

The literature compilation yielded data on 75 unique virus strains, including 51 dsDNA, 1 dsRNA, 7 ssDNA, and 6 ssRNA viruses, and 12 of unknown type (Appendix Table S1). The viruses were isolated from 26 phytoplankton genera, with 38 strains isolated from cyanobacteria, 15 from diatoms, 10 from haptophytes, 7 from chlorophytes, 4 from dinoflagellates, and 1 each from a cryptophyte, raphidophyte, and pelagophyte. The majority was isolated from marine systems (58 vs. 19 from fresh waters). The genome size of the isolates ranges from 4.4 to 560 kb, and capsid diameter ranges from 22 to 310 nm. In our analyses, we consider relationships across all viruses, and also relationships within the dsDNA viruses, which are the most numerous and sometimes show distinct patterns compared to single-stranded viruses.

### Empirical trait relationships

We tested a variety of hypothesized correlations between viral traits, and between viral and host traits. In brief, burst size is most strongly related to the ratio 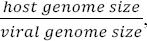 which we refer to as the genome size ratio. Latent period is most strongly related to a combination of host growth rate and the genome size ratio. Figure 1A shows that burst size ranges over four orders of magnitude, and that a larger host genome size, relative to viral genome size, is correlated with a greater burst size. The genome size ratio can explain half of the variation in burst size (R^2^ = 0.49, F_1,44_ = 33, p < 0.001), and the relationship is strongest for dsDNA viruses, with greater variability for the single-stranded viruses that infect diatoms. A similar pattern is found when comparing total viral nucleotide output to host genome size (Fig. S3; R^2^ = 0.54, F_1,5_ = 16, p = 0.01); for dsDNA viruses these quantities tend to be directly proportional, which means that the number of nucleotides in released virions is similar to the number in the host genome.

**Figure 1.**
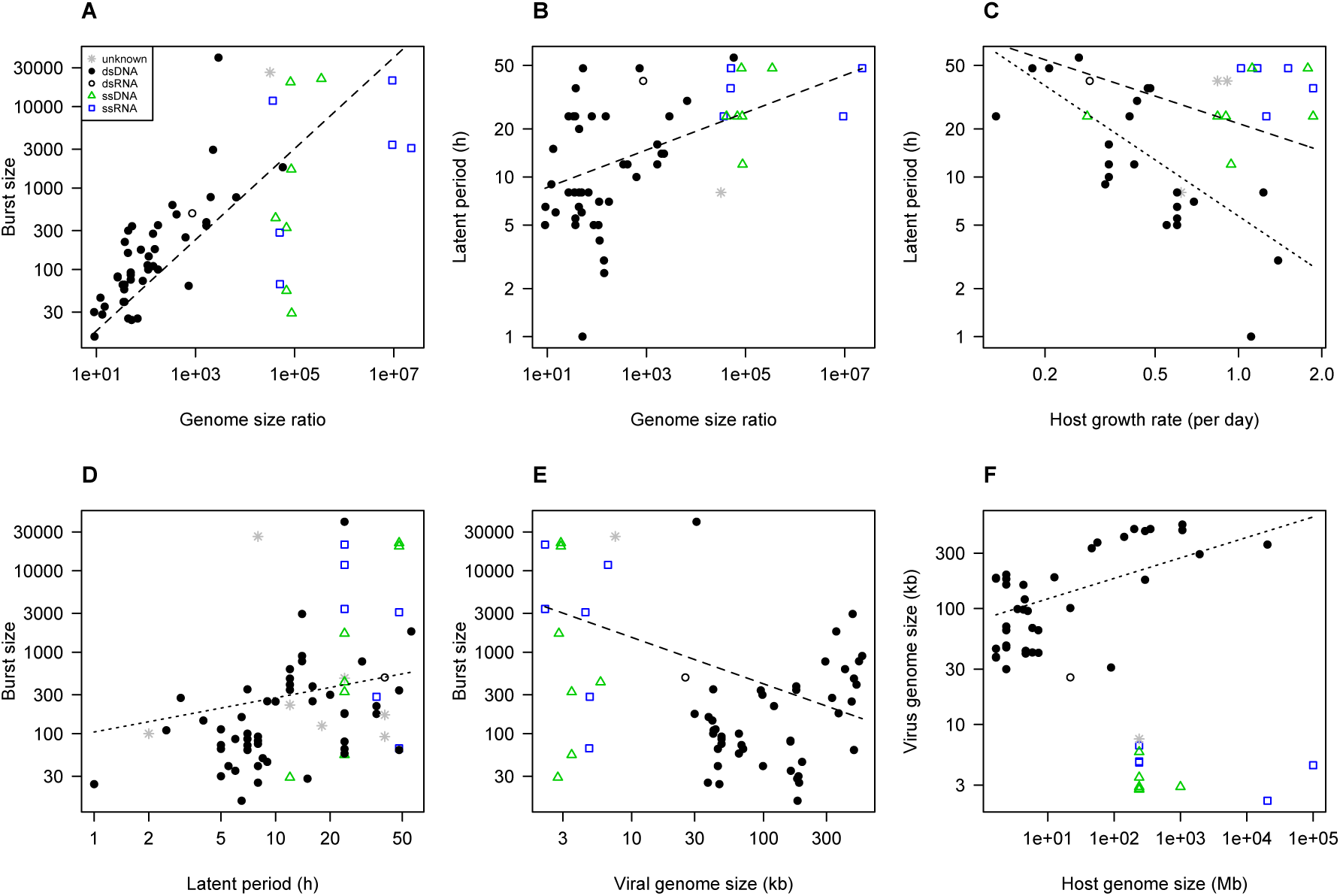
Empirical patterns of trait variation across phytoplankton viruses. (A) Burst size vs. the genome size ratio 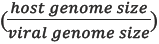. (B) Latent period vs. the genome size ratio. (C) Latent period vs. host growth rate. (D) Burst size vs. latent period. (E) Burst size vs. viral genome size. (F) Virus genome size vs. host genome size. Dashed lines show fitted relationships for all strains, dotted lines show fitted relationships for dsDNA strains. Fitted relationships are from mixed models in which host genus, host taxon, virus type, and publication were included as random effects.

Latent period also increases with the genome size ratio, though less steeply than burst size (Fig. 1B; R^2^ = 0.30, F_1,14_ = 9.3, p = 0.008). For dsDNA viruses there is a stronger relationship between host growth rate and latent period, with latent period declining about 10-fold for a 10-fold increase in host growth rate, and latent period roughly equal to half of the host doubling time (Fig. 1C; R^2^ = 0.47, F_1,9_ = 29, p < 0.001). This pattern is weaker when single-stranded viruses are included (R^2^ = 0.14, F_1,20_ = 6.6, p = 0.02). In a multivariate model, both genome size ratio and host growth rate are significant predictors of latent period, and they jointly explain 38% of variation in latent period across all viruses, and 57% of variation for dsDNA viruses. In contrast, burst size is unrelated to host growth rate (Fig. S3).

Several hypothesized trait correlations are weaker but statistically significant. Burst size and latent period are positively correlated, but only within the dsDNA strains; this corresponds to a 5-fold increase in burst size across a 50-fold increase in latent period (Fig. 1D; R^2^ = 0.05, F_1,22_ = 4.6, p = 0.043). Larger viruses tend to have a smaller burst size when comparing all strains, corresponding to a ∼100-fold decrease in burst size over a 100-fold increase in genome size (Fig. 1E; R^2^ = 0.2, F_1,20_= 11.7, p = 0.003). It is also noteworthy that within the dsDNA viruses there is a trend for larger viruses to infect larger hosts (R^2^ = 0.27, F_1,5.2_ = 5.9, p = 0.058; Fig. 2F), while the smallest viruses, which are single-stranded, have only been isolated from relatively large eukaryotes so far (Fig. 1F). There is no relationship between latent period and viral genome size (Fig. S3).

**Figure 2.**
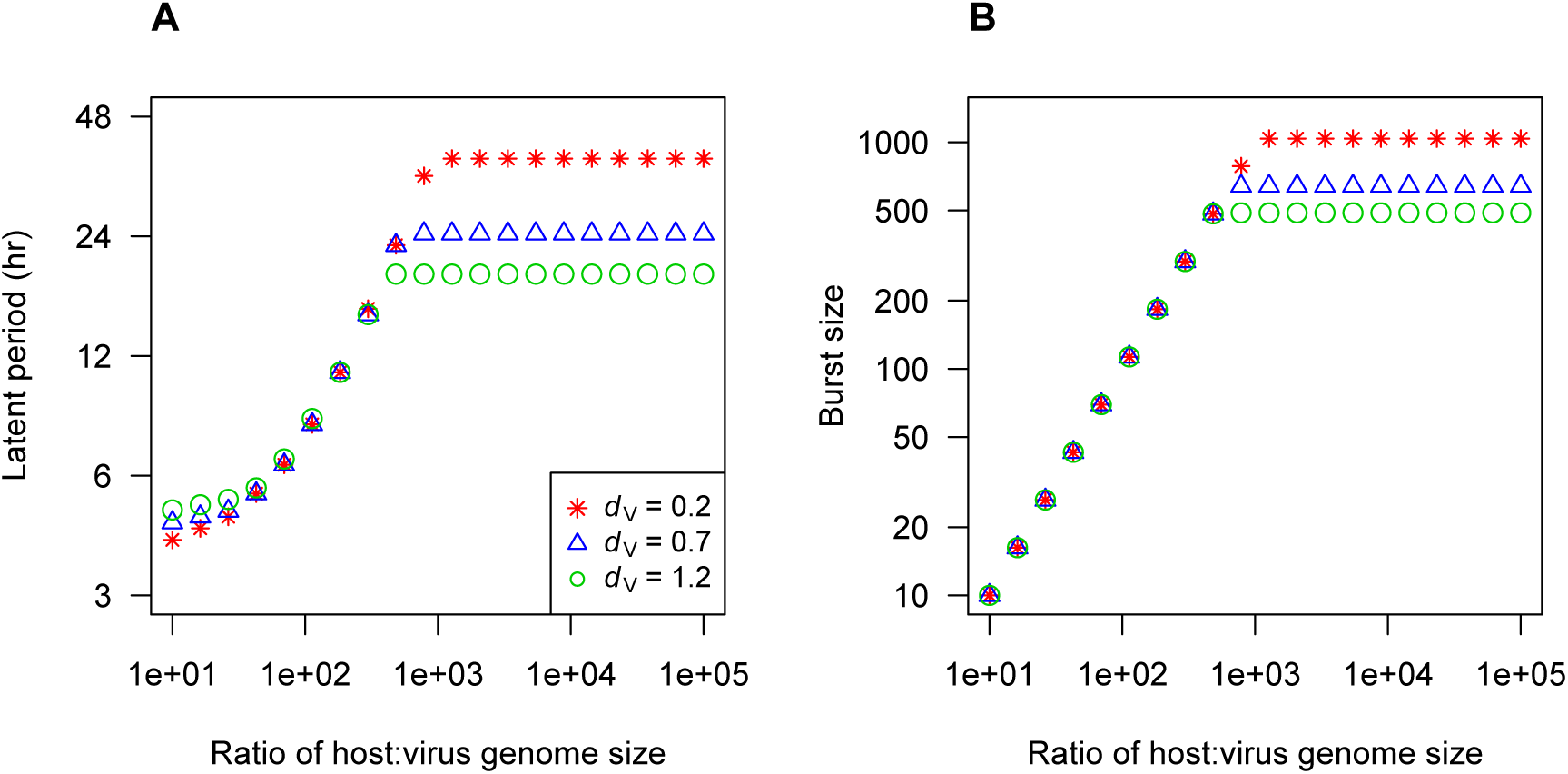
Evolved latent period (A) and burst size (B) as a function of the host:virus genome size ratio. In both panels, the x-axis shows the maximum permitted burst size, which is interpreted here as the ratio of host:virus genome sizes. The y-axis shows the traits of the genotypes that persist in the model. In the model, burst size is calculated instantaneously from the host growth rate (eqns. 5-7); panel (B) shows the burst size for each genotype at maximum host growth, analogous to nutrient-sufficient culture conditions. Results are shown for three virion decay rates (*dv*, d^-1^). *μ*_*max*_ = 1 d^-1^, *r*_*max*_ = 480. virions d^-1^.

### Model of viral trait evolution vs. host genome size

The empirical results suggest that relative genome size and host growth rate are important drivers of viral trait evolution, while burst size and latent period are themselves not strongly correlated across viral strains. We used a model to ask whether these patterns can be explained by mechanistic theory for the evolution of latent period. To ask how host genomic resources might influence viral traits, we used a model in which latent period and burst size can evolve, but burst size is constrained by an upper limit. This limit could have various causes, but here we will imagine that the total number of nucleotides available for viral genome synthesis is approximately equal to the size of the host genome, and therefore the maximum burst size is given by the host:virus genome size ratio (Fig. 1A).

The model predicts a nonlinear relationship between latent period / burst size and the genome size ratio. At low size ratios, selection leads to a strategy that maximizes burst size, i.e. latent period increases until the maximal burst size is reached. This means that both latent period and burst size increase steeply as the genome size ratio increases, until a threshold is reached at a size ratio of ∼1000 (Fig. 2). Above this size ratio, burst size is not maximized and is unrelated to the size ratio. The value of other model parameters affects the asymptotic strategy. For example, a lower virion decay rate selects for longer latent period and greater burst size (Fig. 2). The mortality rate of the host and the concentration of nutrient input have modest effects; reduced host mortality slightly increases asymptotic latent period and burst size, and increased nutrient input slightly reduces latent period (Fig. S4, S5).

These results are consistent with the data showing burst size and latent period tend to increase with the genome size ratio (Fig. 1A-B). They are also consistent with the fact that latent period seems to be capped at about 50 hrs, and may be consistent with the fact that burst size for single-stranded viruses, which infect very large hosts, is highly variable.

### Modeled effects of host growth rate

The model predicts that latent period is inversely proportional to host growth rate, while burst size is insensitive to host growth rate (Fig. 3, Fig. S6). In other words, a higher host growth rate allows the optimal burst size to be achieved with a shorter latent period. We also varied *r*_max_, which accounts for factors beyond host growth rate that influence the rate of virion production. When burst size is maximized, a higher rmax reduces the latent period (Fig. 5); when burst size is not maximized, as observed at high genome size ratios, a higher *r*_max_ increases the burst size instead (Fig. S6). Overall, these results are consistent with the empirical patterns showing a negative correlation between latent period and host growth rate, but no relationship between burst size and growth rate (Fig. 1C, Fig. S6). The results are also consistent with the fact that there is only a weak correlation between burst size and latent period across viruses (Fig. 1D). This is because variation across host species in growth rate or variation across viruses in *r*_max_ will decouple burst size and latent period across viral strains, even though there is a mechanistic connection between the traits.

**Figure 3.**
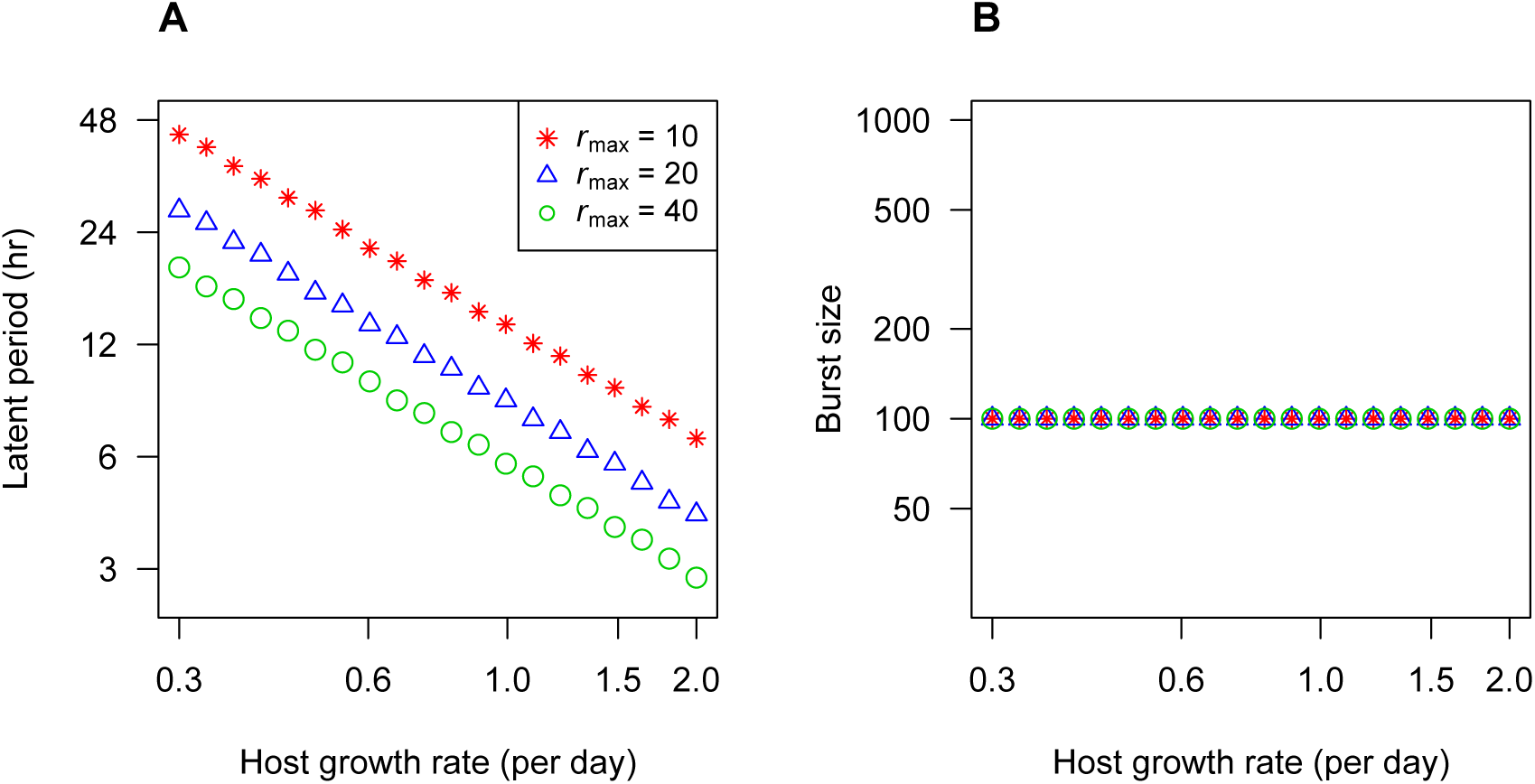
Evolved latent period (A) and burst size (B) as a function of host growth rate. In both panels, the x-axis shows the growth rate of the host, which directly affects the eclipse period and rise rate in the model. The y-axis shows the traits of the genotypes that persist in the model. The maximum burst size (host:virus genome size ratio) is 100 for all cases. Results are shown for three values of *r*_max_ (Eqn. 6), which represents factors beyond host growth rate that affect the virion production rate. *d*_V_ = 0.7 d^-1^

## Discussion

Comparative analyses inevitably deal with noisy patterns, due to the variety of researchers and methods involved. Nonetheless, we find that traits of the phytoplankton host (genome size and growth rate), in combination with virus genome size, can collectively explain ∼40-50% of variation in burst size and latent period across phytoplankton viruses characterized thus far. We also find that a model of viral trait evolution, parameterized with realistic values, produces patterns similar to the empirical results, which lends support to the hypothesized underlying mechanisms. Our interpretation of these results is that phytoplankton cells are a sparse resource in a world that is relatively hazardous. This leads to selection for latent periods that are long enough to exhaust host resources before lysis, for viruses that are not extremely small compared to their host. This results in a correlation between total nucleotide output and host genome size, or between burst size and the host:virus genome size ratio. In addition, latent period is jointly influenced by the genome size ratio and the host growth rate, because the physiology of more rapid host growth allows for more rapid virion production. When the genome size ratio is very large (>∼1000), it may be that latent period would have to be several days long in order to exhaust host resources, which increases the likelihood of host mortality during infection. For these cases, other factors may determine the evolution of latent period, such as mortality rates of the host and free virions. In total, our results argue that a trait-based approach to viral ecology is promising, and important aspects of community dynamics and evolution may be predictable from a relatively simple set of underlying principles.

The patterns across viral strains can be compared with short-term experiments on a single strain, where host growth is manipulated and virus reproduction is characterized. Experiments with *E. coli* typically show that increased host growth rate decreases latent period and increases burst size (You et al. 2002). In an experiment with *Synechococcus*, an increase in growth due to stirring did not alter latent period (Wilson et al. 1996), while experiments with *Micromonas* and *Phaeocystis* found that both nitrogen- and phosphorus-limited growth increased latent period and reduced burst size (Maat and Brussaard 2016). Observed correlations between burst size and host growth rate differ from our empirical and model results (Fig. 3, Fig. S3). In this context, it is important to note that short-term plastic responses of a strain may differ from the evolved strategies that vary across strains and hosts. Indeed, in our model the immediate effect of faster host growth is to increase the rate of viral production, which will lead to a greater burst size for a particular latent period. However, over the long term the optimal genotype is one that reduces the latent period while maintaining the same burst size.

Our compilation is relevant to some additional questions not addressed by our model, which focused on the evolution of burst size and latent period. Viruses of phytoplankton vary greatly in size, and the maintenance of this size variation remains to be explained. All else equal, larger virions should take longer to synthesize/assemble, and if host resources are limiting then fewer virions can be produced. Both considerations should reduce burst size for larger viruses, but this is only modestly evident in the data (Fig. 1E). A similar pattern is found when (burst size)/(latent period) is used to approximate the rate of virion production (Fig. S7). There is evidence that larger viral genome size allows for better control of host metabolism, which could increase the rate of viral production and/or the contribution of *de novo* nucleotide synthesis (Lindell et al. 2005, Hurwitz et al. 2013). Modeling a general mechanistic relationship between genome size and viral reproduction will require a complex model of host-viral metabolism and its evolution (Bragg and Chisholm 2008, Birch et al. 2012).

The patterns in our compilation are most evident for dsDNA viruses, which have been studied in much greater detail than other types. It is possible that single-stranded viruses are under distinct selective pressures as a consequence of their very small size or different type of interaction with host systems of transcription/translation/replication. In addition, at this point single-stranded viruses of phytoplankton have been isolated primarily from *Chaetoceros* spp. (e.g., Nagasaki et al. 2005, Tomaru et al. 2008, Kimura and Tomaru 2015). The authors of those studies further note that burst size is hard to define, because it appears that new viral particles are released prior to lysis. Therefore, a somewhat different mode of infection may be part of the reason the strains show different patterns in our analyses. An additional factor that could add noise or bias to the data is the fact that viral strains maintained in culture could evolve in response to propagation, during which they are periodically exposed to relatively high host density, which could select for a shorter latent period.

The results of this study can contribute to models of virus-microbe interactions, community structure, and ecosystem consequences. Recent models have explored how viruses can promote host diversity (Thingstad et al. 2014), how hosts and viruses with overlapping host ranges can coexist (Jover et al. 2013), and how viruses alter the dynamics of standard ecosystem models (Weitz et al. 2015), among other topics. The relationships between virus and host traits described here could be used to constrain model parameters, allowing host and virus community structure to self-organize across environmental gradients (Follows and Dutkiewicz 2011). Cell size is a ‘master trait’ that influences many aspects of plankton ecology (Finkel et al. 2009, Edwards et al. 2012), and size has been used to structure trait-based models via allometric scaling relationships (Dutkiewicz et al. 2011). Genome size is well-correlated with cell size in phytoplankton and other organisms (Veldhuis et al. 1997), and therefore the results of the current study could be used to derive scaling relationships and incorporate viral dynamics into size-structured models of plankton ecosystems. Successful development of virus community models will also require considerable empirical progress on how host range and host resistance coevolve, and how infection network structure is related to both viral traits (burst size, adsorption rate, etc.) and host traits (resistance and its physiological costs).

## References

Abedon, S. T., P. Hyman, and C. Thomas. 2003. Experimental Examination of Bacteriophage Latent-Period Evolution as a Response to Bacterial Availability Experimental Examination of Bacteriophage Latent-Period Evolution as a Response to Bacterial Availability. Applied and Environmental Microbiology 69:7499–7506.

Bartón, K. 2016. MuMIn: Multi-Model Inference. R package version 1.15.6.

Bates, D., Maechler, M., Bolker, B., and S. Walker. 2015. Fitting Linear Mixed-Effects Models Using lme4. Journal of Statistical Software 67:1–48.

Birch, E. W., N. A. Ruggero, and M. W. Covert. 2012. Determining Host Metabolic Limitations on Viral Replication via Integrated Modeling and Experimental Perturbation. PLoS Computational Biology 8.

Bonachela, J. A., and S. A. Levin. 2014. Evolutionary comparison between viral lysis rate and latent period. Journal of Theoretical Biology 345:32–42.

Bragg, J. G., and S. W. Chisholm. 2008. Modeling the fitness consequences of a cyanophage-encoded photosynthesis gene. PLoS ONE 3:1–9.

Breitbart, M. 2012. Marine Viruses: Truth or Dare. Annual Review of Marine Science 4:425–448.

Brown, C. M., J. E. Lawrence, and D. A. Campbell. 2006. Are phytoplankton population density maxima predictable through analysis of host and viral genomic DNA content? Journal of the Marine Biological Association of the UK 86:491.

Brown, C. M., D. A. Campbell, and J. E. Lawrence. 2007. Resource dynamics during infection of Micromonas pusilla by virus MpV-Sp1. Environmental Microbiology 9:2720–2727.

Brussaard, C. P.D. 2004. Viral control of phytoplankton populations: a review. The Journal of Eukaryotic Microbiology 51:125–138.

Daines, S. J., J. R. Clark, and T. M. Lenton. 2014. Multiple environmental controls on phytoplankton growth strategies determine adaptive responses of the N:P ratio. Ecology Letters 17:414–425.

Edwards, K. F., M. K. Thomas, C. A. Klausmeier, and E. Litchman. 2012. Allometric scaling and taxonomic variation in nutrient utilization traits and maximum growth rate of phytoplankton. Limnology and Oceanography 57:554–566.

Finkel, Z. V., J. Beardall, K. J. Flynn, A. Quigg, T. A. V Rees, and J. A. Raven. 2009. Phytoplankton in a changing world: Cell size and elemental stoichiometry. Journal of Plankton Research 32:119–137.

Fischer, M. G. 2016. ScienceDirect Giant viruses come of age. Curr Opin Microbiol 31:50–57.

Gudelj, I., J. S. Weitz, T. Ferenci, M. Claire Horner-Devine, C. J. Marx, J. R. Meyer, and S. E. Forde. 2010. An integrative approach to understanding microbial diversity: from intracellular mechanisms to community structure. Ecology Letters 13:1073–84.

Hurwitz, B. L., S. J. Hallam, and M. B. Sullivan. 2013. Metabolic reprogramming by viruses in the sunlit and dark ocean. Genome biology 14:R123.

Kimura, K., and Y. Tomaru. 2015. Discovery of two novel viruses expands the diversity of single-stranded DNA and single-stranded RNA viruses infecting a cosmopolitan marine diatom. Applied and Environmental Microbiology 81:1120–1131.

Koonin E. V., and V. V. Dolja. 2013. A virocentric perspective on the evolution of life. Current Opinion in Virology 3:546–557.

Kuznetsova, A., P. B. Brockhoff, and R. H.B. Christensen. 2016. lmerTest: Tests in linear mixed effects models. R package version 2.0-32.

Lang A. S., M. L. Rise, A. I. Culley, and G. Steward. 2009. RNA viruses in the sea. FEMS Microbiol Rev 33:295–323.

Levin, B., F. Stewart, and L. Chao. 1977. Resource-limited growth, competition, and predation: a model and experimental studies with bacteria and bacteriophage. American Naturalist 111:3–24.

Lindell, D., J. D. Jaffe, Z. I. Johnson, G. M. Church, and S. W. Chisholm. 2005. Photosynthesis genes in marine viruses yield proteins during host infection. Nature 438:86–9.

Litchman, E., and C. A. Klausmeier. 2008. Trait-Based Community Ecology of Phytoplankton. Annual Review of Ecology, Evolution, and Systematics 39:615–639.

Maat, D. S., and C. P.D. Brussaard. 2016. Both phosphorus- and nitrogen limitation constrain viral proliferation in marine phytoplankton. Aquatic Microbial Ecology 77:87–97.

Murray, A. G., and G. A. Jackson. 1992. Viral Dynamics A Model of the Effects of Size Shape Motion and Abundance of Single-Celled Planktonic Organisms and Other Particles. Marine ecology progress series 89:103–116.

Nagasaki, K., Y. Tomaru, Y. Takao, K. Nishida, Y. Shirai, H. Suzuki, et al. 2005. Previously unknown virus infects marine diatom. Applied and Environmental Microbiology 71:3528–3535.

Rosario, K., S. Duffy, and M. Breitbart. 2012. A field guide to eukaryotic circular single-stranded DNA viruses: insights gained from metagenomics. Arch Virol 157:1851–1871.

Soetaert, K. T. Petzoldt, and R. W. Setzer. 2010. Solving Differential Equations in R: Package deSolve. Journal of Statistical Software, 33:1–25.

Steward, G. F., A. I. Culley, and E. M. Wood-Charlson. 2013. Marine Viruses. Pages 127–144 In: Levin, S.A. (ed.) Encyclopedia of Biodiversity. Second Edition.

Suttle, C. A. 2007. Marine viruses—major players in the global ecosystem. Nature Reviews Microbiology 5:801–12.

Thomas, R., N. Grimsley, M.-L. Escande, L. Subirana, E. Derelle, and H. Moreau. 2011. Acquisition and maintenance of resistance to viruses in eukaryotic phytoplankton populations. Environmental microbiology 13:1412–20.

Tomaru, Y., Y. Shirai, H. Suzuki, T. Nagumo, and K. Nagasaki. 2008. Isolation and characterization of a new single-stranded DNA virus infecting the cosmopolitan marine diatom Chaetoceros debilis. Aquatic Microbial Ecology 50:103–112.

Tummers, B. DataThief III. 2006 <http://datathief.org/>

Van Etten, J. L., D. E. Burbank, J. Joshi, and R. H. Meints. 1984. DNA synthesis in a Chlorella-like alga following infection with the virus PBCV-1. Virology 134:443–449.

Veldhuis, M. J. W., T. L. Cucci, and M. E. Sieracki. 1997. Cellular DNA content of marine phytoplankton using two new fluorochromes: taxanomic and ecological implications. Journal of Phycology 33;527–541.

Villarreal L. P., and G. Witzany. 2010. Viruses are essential agents within the roots and stem of the tree of life. Journal of Theoretical Biology 262:698–710.

Wang, I.-N., D. E. Dykhuizen, and L. B. Slobodkin. 1996. The evolution of phage lysis timing. Evolutionary Ecology 10:545–558.

Weitz, J. S., C. A. Stock, S. W. Wilhelm, L. Bourouiba, M. L. Coleman, A. Buchan, et al. 2015. A multitrophic model to quantify the effects of marine viruses on microbial food webs and ecosystem processes. The ISME Journal 9:1352–1364.

Westoby, M., and I. J. Wright. 2006. Land-plant ecology on the basis of functional traits. Trends in Ecology & Evolution 21:261–268.

Wikner, J., J. J. Vallino, G. F. Steward, D. C. Smith, and F. Azam. 1993. Nucleic-Acids from the Host Bacterium as a Major Source of Nucleotides for 3 Marine Bacteriophages. Fems Microbiology Ecology 12:237–248.

Wilson, W. H., N. G. Carr, and N. H. Mann. 1996. The effect of phosphate status on the kinetics of cyanophage infection in the oceanic cyanobacterium Synechococcus sp. WH7803. Journal of Phycology 32:506–516.

You, L., P. F. Suthers, and J. Yin. 2002. Effects of Escherichia coli physiology on growth of phage T7 in vivo and in silico. Journal of Bacteriology 184:1497–1500.

